# Evolutionary origin of prolonged delayed fertilization in the Fagaceae

**DOI:** 10.64898/2026.01.15.699616

**Authors:** Takenori Shagawa, Chihiro Myotoishi, Tetsukazu Yahara, Ryosuke Imai, Min Deng, Akiko Satake

## Abstract

The temporal organization of flowering, fertilization, and fruiting is a fundamental axis of life-history evolution in angiosperms. While most species complete fruit development within a single growing season (“1-year fruiting”), many Fagaceae species delay fertilization and fruit maturation until the year following flowering (“2-year fruiting”). Despite its ecological prevalence, the evolutionary origins of this strategy and its coordination with other functional traits remain poorly understood. In this study, we investigated the evolutionary origins of the two-year fruiting strategy by reconstructing ancestral states on a phylogeny comprising 88 species that represent all eight genera of Fagaceae. We further employed phylogenetic comparative analyses to test whether the fruiting trait is evolutionarily constrained by pollination mode (animal versus wind-pollinated) or leaf habit (evergreen versus deciduous). Our results support a single origin of 2-year fruiting in the common ancestor of the major clade excluding *Fagus* and *Trigonobalanus*, followed by multiple independent reversions to 1-year fruiting in lineages including *Castanea*, *Quercus*, and *Castanopsis*. Ancestral-state reconstructions of pollination mode and leaf habit strongly support entomophily and an evergreen leaf habit as ancestral traits of Quercoideae, predating the emergence of the 2-year fruiting strategy. Although entomophily and evergreen habit were ancestral in Quercoideae, our correlation analyses indicate that transitions in fruiting strategy did not depend on changes in these traits. These results provide macroevolutionary evidence that extreme extensions of the interval between pollination and fertilization can be evolutionarily stable yet rarely re-evolve once lost over long evolutionary timescales.

## 1 Introduction

The temporal organization of key reproductive events—flowering, fertilization, and fruit maturation—plays a fundamental role in shaping organismal fitness and underlies phenological and life-history diversification in angiosperms (Schwarz 2003; Satake et al. 2022). Most angiosperms complete the transition from pollination to fertilization within 24–48 hours (Williams 2008; Williams and Reese 2019). However, a subset of taxa has evolved reproductive strategies in which fertilization is substantially delayed, ranging from several days to more than a year after pollination (Sogo and Tobe 2006; Williams 2008; Williams and Reese 2019). These strategies, collectively referred to as delayed fertilization (Sogo and Tobe 2006; Williams 2008; Williams and Reese 2019), represent a striking deviation from the canonical angiosperm reproductive phenology and provide a unique opportunity to explore the adaptive significance of extended reproductive cycles in plants (Schoonderwoerd and Friedman 2016; Satake and Kelly 2021; Deng et al. 2022; Satake et al. 2023; Yao et al. 2023; Shagawa et al. 2025).

The Fagaceae, including oaks and beeches, present a particularly striking example of such deviation. In many Fagaceae species, fertilization is delayed until the year following anthesis, resulting in a “2-year fruiting” strategy in which fertilization and subsequent fruit maturation occur in the year following flowering. In contrast, under the “1-year fruiting” strategy, the entire reproductive sequence—from anthesis through fertilization to fruit maturation—occurs within a single growing season (Fig. 1; Benson 1894; Sogo and Tobe 2006; Deng et al. 2022). Across the family, 1-year and 2-year fruiting strategies coexist, with their prevalence differing markedly among genera: all *Fagus* species show 1-year fruiting, whereas the majority of *Castanopsis* and *Lithocarpus* species exhibit 2-year fruiting, and *Quercus* shows an intermediate pattern (Satake & Kelly 2021; Araye et al. 2023). Although the 2-year fruiting with prolonged delayed fertilization has been recognized for more than a century, its evolutionary origins, the frequency and direction of transitions between 1-year and 2-year fruiting, and their adaptive or ecological significance remain poorly understood (Satake and Kelly 2021; Satake et al. 2023; Shagawa et al. 2025).

**FIGURE 1.**
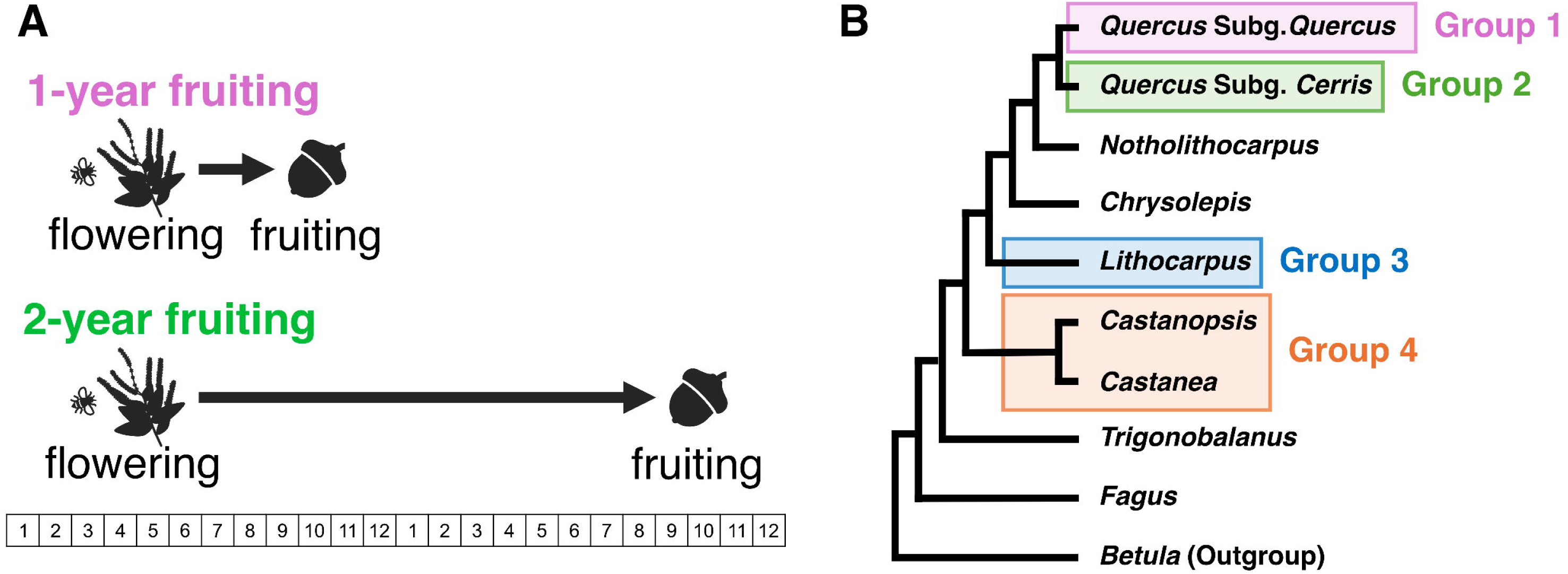
Fruiting strategies in Fagaceae and ancestral state reconstruction of fruiting traits. **(A)** 1-year and 2-year fruiting strategies observed in Fagaceae. In the 1-year fruiting species, flowering and fruit maturation are completed within a single year, whereas in the 2-year fruiting species, fruit maturation occurs in the year following flowering. **(B)** Phylogenetic relationships among Fagaceae genera and the grouping used for ancestral state reconstruction of the fruiting traits. Four groups were defined for the analysis: Group 1 (*Quercus* Subg. *Quercus*), Group 2 (*Quercus* Subg. *Cerris*), Group 3 (*Lithocarpus*), and Group 4 (*Castanopsis* and *Castanea*). The model assumes different transition rates among these groups. Ancestral state reconstruction was performed for the 52 alternative model settings shown in Figure A2, and the optimal model was selected based on the marginal likelihood values calculated for each model.

To address this gap, we integrated phylogenetic comparative analyses with a comprehensive fruiting-trait dataset from 88 species spanning all eight genera of Fagaceae to reconstruct the evolutionary history of prolonged delayed fertilization underlying the 2-year fruiting strategy. We further tested for correlated evolution between fruiting strategy and two traits hypothesized to constrain it: pollination mode and leaf habit. Satake and Kelly (2021) suggested that delayed fertilization is more likely to evolve in animal-pollinated species than in wind-pollinated species, owing to stronger pollen competition and interspecific interference under seasonal environments. Leaf habit may provide a second, independent axis of constraint. Evergreen species, capable of year-round photosynthesis, may sustain the prolonged development and overwinter maintenance of ovules or young fruits (Givnish et al. 2002). In contrast, deciduous species—with shorter periods of carbon assimilation and greater risk of carrying immature fruits through winter—may be constrained to complete reproduction within a single growing season.

Here, we use phylogenetic comparative methods to test for correlated evolution among fruiting strategy, pollination mode, and leaf habit across the Fagaceae. Together, these analyses provide a framework for reconstructing the evolutionary history of 2-year fruiting in the Fagaceae, including its timing and frequency of origin and loss, and for evaluating whether shifts in fruiting strategy have been evolutionarily coordinated with changes in pollination mode or leaf habit.

## 2 Materials and methods

### 2.1 Trait data collection

We followed the taxon-sampling scheme of Zhou et al. (2022), which includes 90 Fagaceae species across eight genera and provides a genome-scale nuclear phylogeny. To maintain consistency with recent molecular phylogenetic frameworks (Hipp et al. 2020; Zhou et al. 2022; Liu et al. 2025; Shen et al. 2025; Yang et al. 2025), we adopted updated generic circumscriptions: species listed as *Cyclobalanopsis* and *Formanodendron* in the eFlora of China were treated as *Quercus* and *Trigonobalanus*, respectively (Brach and Song 2006), and species listed as *Lithocarpus* in the eFlora of North America were treated as *Notholithocarpus* (Nixon 1997). We excluded *Quercus liaotungensis* Koidz. as it is treated as a synonym of *Q. mongolica* Fisch. ex Ledeb. in eFlora of China (Brach and Song 2006), which was already included in the dataset.

We compiled fruiting trait data–1-year or 2-year fruiting–from the eFlora of China (Brach & Song 2006) and the eFlora of North America (Nixon 1997) (Table S1), for 88 species encompassing all eight genera. For 19 species for which information on the three traits was unavailable or potentially inaccurate in the eFlora of China or the eFlora of North America, we consulted additional references (listed in Table S1). For eight species for which information was unavailable from the literature, we examined online herbarium specimens from the herbarium databases listed in Table S2. In contrast, *Lithocarpus tephrocarpus* was classified as a 2-year fruiting species based on field observations. In addition, traits for pollination mode (anemophily or entomophily) and leaf habit (deciduous or evergreen) were also compiled based on the references listed in Table S1. In this study, leafing traits were classified into two categories: deciduous and evergreen. However, some species have been reported to exhibit an intermediate condition, often referred to as “subevergreen” (or semi-deciduous). In such cases, assigning three states to a single trait would prevent the implementation of discrete models in BayesTraits (Pagel et al. 2004; Pagel & Meade 2006). Therefore, we consulted multiple references and assigned each species to either deciduous or evergreen.

Using specimen images from online herbarium databases, we first confirmed species identities based on diagnostic morphological traits and then determined whether each species exhibits 1-year or 2-year fruiting. For each specimen, we examined twigs bearing female reproductive organs. Current-year and previous-year twigs were distinguished using leaf scars as morphological markers. To classify fruiting traits, we treated all duplicate specimens derived from the same collection event as a single specimen group. A species was classified as 2-year fruiting if at least one specimen group contained two distinct developmental stages of female reproductive organs: (i) flowers or fruits retained from the previous year and (ii) flowers produced in the collection year. Species that did not meet this criterion were classified as 1-year fruiting. Because individual herbarium sheets often include only a single developmental stage, evaluating specimen groups rather than individual sheets enabled us to detect cases in which two stages occurred among duplicates from the same collection event. We assumed that flowers originating from a single flowering event develop within the same season; therefore, the coexistence of two clearly distinct developmental stages within a specimen group was interpreted as evidence for 2-year fruiting. *Trigonobalanus excelsa* Lozano, Hern.Cam. & Henao for which reliable information could not be obtained, even after examination of herbarium specimens, was excluded from subsequent analyses.

### 2.2 Phylogenetic reconstruction

We conducted a Bayesian phylogenetic analysis using MrBayes v3.2.7 (Ronquist et al. 2012) using nucleotide alignments of 2,124 nuclear genes provided by Zhou et al. (2022). Before analysis, sequences not belonging to our focal taxa (Table S1) were excluded. We ran four independent Markov chain Monte Carlo (MCMC) analyses, each for 1,000,000 generations with sampling every 500 generations. The first 25% of sampled trees from each run were discarded as burn-in. The remaining post-burn-in trees were pooled to generate a 50% majority-rule consensus tree and to calculate posterior probabilities for nodes (Fig. S1).

The model settings followed Zhou et al. (2022), adopting the optimal partitioning scheme and nucleotide substitution models using PartitionFinder 2 v1.1 (Lanfear et al. 2017) under the Akaike Information Criterion (AIC) (Akaike 1974). To account for topological uncertainty in subsequent ancestral character state reconstructions, we randomly resampled 500 trees from the post-burn-in trees.

### 2.3 Ancestral state reconstruction

We used BayesTraits v3.0.5 (Pagel et al. 2004; Pagel & Meade 2006) to infer the ancestral states of fruiting types, pollination modes, and leaf habits, and to test for correlated evolution among these traits. Individual traits were analyzed under the multistate model, while pairwise correlated evolution between binary traits was tested with the discrete model. To satisfy the requirements of the discrete model, we encoded traits as follows: the fruiting trait (1-year fruiting = 0, 2-year fruiting = 1), the pollination mode (anemophily = 0, entomophily = 1), and the leafing trait (deciduous = 0, evergreen = 1) (see Table S1).

Within the multistate model framework, we compared an all-rates-different model (ARD), which estimates distinct transition rates among all states, with an equal-rates model (ER) that constrains all transitions to be identical, using the Restrict function in BayesTraits. For fruiting traits, we additionally fitted clade-heterogeneous variants in which transition rates were allowed to differ among major clades, given substantial genus-level variations in the prevalence of fruiting traits (Satake & Kelly 2021). To test this, we used the AddPattern function in BayesTraits to assign clade-specific transition rates to four specie-rich clades (each ≥10 species): *Quercus* subg. *Quercus*, *Quercus* subg. *Cerris*, *Lithocarpus*, and *Castanea* + *Castanopsis* (Fig. 1b). This resulted in 52 alternative model specifications (Fig. S2). Furthermore, several *Quercus* species exhibit intraspecific– or even intra–individual polymorphism in fruiting type (e.g., *Q. suber* L.: Elena-Rosselló et al. 1993; Díaz-Fernández et al. 2004). For *Q. suber*, we therefore conducted ancestral state reconstructions under two alternative assumptions, treating the species as 1-year or 2-year fruiting; these were analyzed as separate datasets (“Fruiting_A” and “Fruiting_B”; Table S4). For pollination mode and leaf habit, we compared only the equal-rates (ER) and all-rates-different (ARD) models (i.e., no clade-heterogeneous variants).

In the discrete model (Pagel 1994), we compared four alternative models of trait evolution: (i) an independent ARD model (4 parameters), (ii) an independent ER model (2 parameters), (iii) a dependent-ARE model (8 parameters), and (iv) a dependent-ER model (4 parameters) (Fig. S3). In the independent models, the two binary traits evolve on separate Markov chains, with no effect on each other. In the dependent models, the transition rates for one trait can vary conditionally on the state of the other.

Model selection was performed using Bayes factors (BF) calculated from marginal likelihoods of competing models (Kass & Raftery 1995). We computed 2ln *BF*, defined as 2ln *BF* = 2[ln *m*(*complex*) - ln *m*(*simple*)], where m denotes the marginal likelihood. The resulting 2ln *BF* values were interpreted following Kass and Raftery (1995): positive (2–6), strong (6–10), and very strong (> 10) evidence in favor of the more complex model. We report 2 ln BF and interpret the values following Kass and Raftery (1995) (positive: 2–6; strong: 6–10; very strong: >10).

All BayesTraits analyses were run for 1,250,000 iterations, with the first 250,000 discarded as burn-in and sampling performed every 1,000 iterations. At each step of the MCMC runs, one tree was randomly chosen from a posterior sample of 500 resampled post-burn-in trees to integrate topological and branch-length uncertainty. Marginal likelihoods were estimated using the stepping-stone sampler (Xie et al. 2011) with 100 stones and 10,000 iterations per stone; model-specific values are reported in Table S4. We placed an exponential hyperprior on transition rate parameters, with the mean drawn from a uniform distribution between 0 and 100, and used reversible-jump MCMC to traverse model space (Pagel et al. 2006). Posterior state probabilities were then mapped onto a 50% majority-rule consensus tree that we time-scaled with eight fossil calibrations (age ranges following Zhou et al. (2022)) using the chronos function in the R package ape (Paradis et al. 2004; Paradis & Schliep 2019).

## 3 Results

### 3.1 Ancestral trait inference supports a two-year fruiting origin

The best-fit model for ancestral state reconstruction for the fruiting trait was the single transition rate model, in which transitions between one-year and two-year fruiting occurred at equal rates across the tree (0.757 ± 0.066; Fig. 2). Under this model, node N3—uniting the six clades except *Fagus* and *Trigonobalanus*—was strongly supported as the 2-year fruiting (86.8 ± 2.56%; Fig. 2). In contrast, the ancestral state of the deeper nodes N1 and N2, corresponding to the early divergences involving *Fagus* and *Trigonobalanus*, could not be confidently resolved (Fig. 2), reflecting limited resolution at the base of the lineage.

**FIGURE 2.**
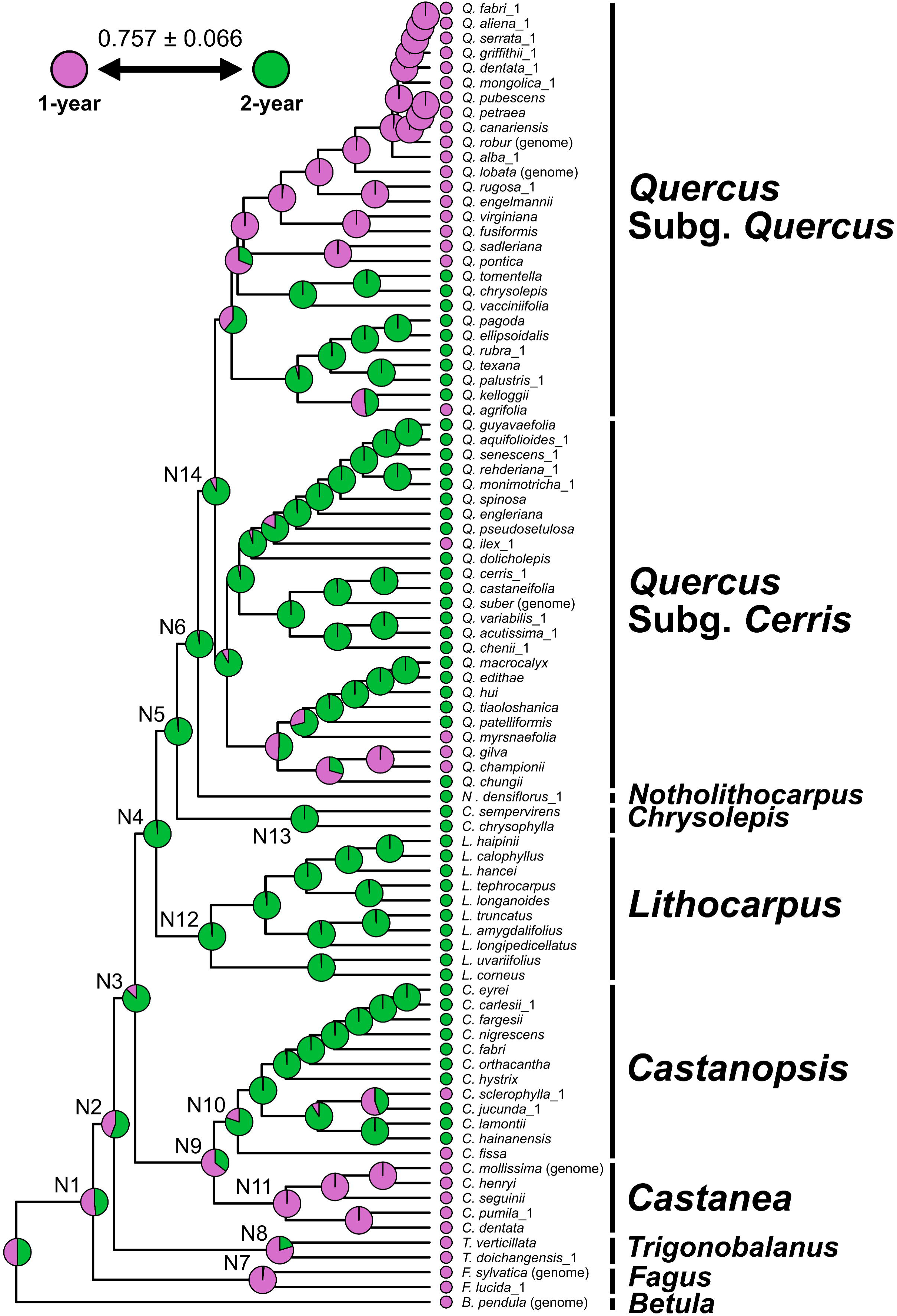
Estimated evolutionary history of fruiting traits based on the best-supported result from the multistate model analyses when *Quercus suber* is treated as a 2-year fruiting. Pink and green represent 1-year and 2-year fruiting, respectively. The pie chart at each node within the phylogenetic tree represents the proportional probability of 1-year and 2-year fruiting. The value below the phylogenetic tree indicates the relative transition rate between the 1-year and 2-year fruiting.

The 2-year fruiting state was consistently retained after the divergence of *Lithocarpus*, *Chrysolepis*, and *Notholithocarpus* at node N4-6, each supported by posterior probabilities exceeding 95% (Fig. 2). Within the clade of *Castanea* and *Castanopsis*, the ancestral node N9 was equivocal, with a posterior probability of 64.3 ± 4.76% marginally supporting the 1-year fruiting. In contrast, the common ancestors of all other genera containing the 2-year fruiting species were inferred to be the 2-year fruiting, with strong posterior support (*Castanopsis*: 80.2 ± 1.52% at node N10; *Lithocarpus*: 98.7 ± 1.06% at “N12”; *Chrysolepis*: 99.9 ± 0.0702% at node N13; *Quercus*: 92.6 ± 4.45% at node N14). Conversely, the ancestors of genera containing only the 1-year fruiting were inferred to be the 1-year fruiting (*Fagus*: 98.3 ± 1.12% at node N7; *Trigonobalanus*: 79.6 ± 8.19% at node N8; *Castanea*: 99.2 ± 0.602% at node N11).

Our analyses also showed that transitions from the 2-year to the 1-year fruiting occurred multiple times independently across lineages. The 1-year fruiting species arose predominantly from the 2-year fruiting clades in *Castanopsis*, *Quercus* subg. *Cerris* and subg. *Quercus*. Notably, the common ancestor of Sect. *Ponticae*, Sect. *Virentes*, and Sect. *Quercus* within subg. *Quercus* was reconstructed with a posterior probability of 99.2 ± 0.644% for the 1-year fruiting, consistent with the shared state across these sections. In addition, the analysis treating *Quercus suber* as a 1-year fruiting species yielded results consistent with those described above (Fig. S4a).

### 3.2 Estimated evolutionary history of the pollination and leafing traits

Independent analyses for pollination mode and leaf habit showed that, for both traits, models constraining transition rates were selected, and the estimated relative transition rates were 0.200 ± 0.116 and 0.572 ± 0.226, respectively (Fig. S4B, C). For pollination mode, the node N2 comprising *Trigonobalanus* and its sister clade was strongly supported as entomophilous with a posterior probability of 96.3 ± 2.46%, although the ancestral state of Fagaceae was ambiguous (Fig. S4B). This state was retained until the divergence of *Lithocarpus*, after which the common ancestor of *Notholithocarpus* and *Quercus* at the node N6 was strongly supported as anemophilous (88.9 ± 4.13%). Independent origins of anemophily were further supported in *Fagus*, *Trigonobalanus doichangensis*, and *Quercus* (Fig. S4B).

For leaf habit, the ancestral state of Fagaceae was likewise ambiguous, as indicated by node N1, which showed comparable likelihoods for deciduous and evergreen states (Fig. S4C). However, the node N2 comprising *Trigonobalanus* and its sister clade was strongly supported as evergreen (94.5 ± 4.70%). The common ancestors of the evergreen genera were supported as evergreen with posterior probabilities exceeding 85.9% (*Trigonobalanus*: 85.9 ± 7.59% at node N8; *Castanopsis*: 98.7 ± 1.07% at node N10; *Lithocarpus*: 99.3 ± 0.716% at node N12; *Chrysolepis*: 99.9 ± 0.0507% at N13; *Quercus*: 97.5 ± 1.68% at node N14; Fig. S4C), indicating that deciduousness observed in *Castanea* and some lineages of *Quercus* is a derived condition. Because the ancestral state of Fagaceae remains unresolved, it is not possible to determine whether the deciduous habit of Fagus represents an ancestral or derived state.

### 3.3 Testing correlated evolution between two traits

Finally, we examined whether fruiting traits evolved in correlation with pollination or leafing traits. The independent ER model was selected as the best-fitting model for both traits (Fig. 3), suggesting that there is no support for correlated trait evolution. In both analyses, transitions in fruiting, pollination, and leafing traits occurred independently, with estimated transition rates closely matching those obtained when ancestral states were reconstructed independently for each trait (Figs. 2, 3, S4).

**FIGURE 3.**
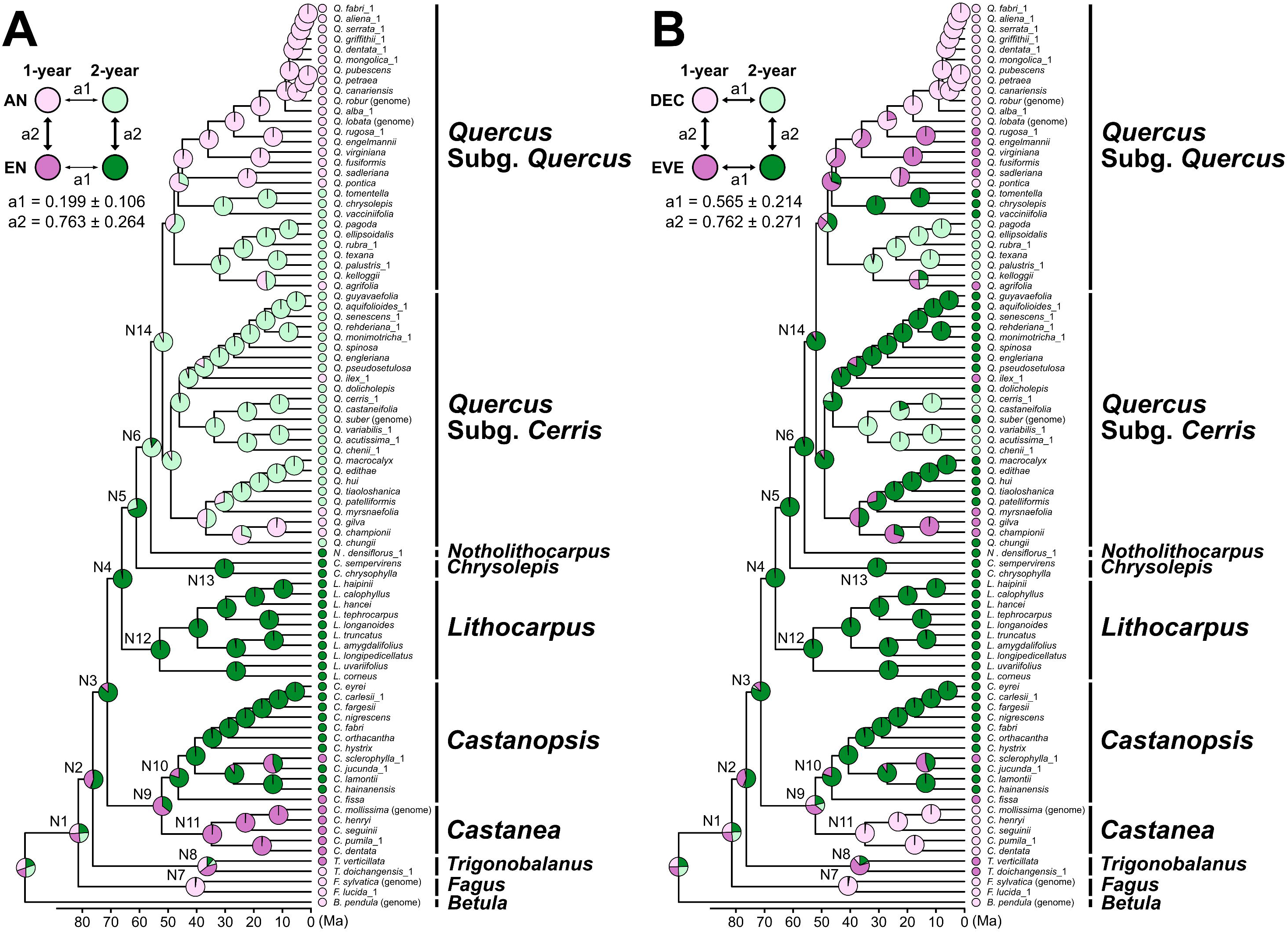
Estimated evolutionary history of the correlated evolution between fruiting trait and (A) pollination trait and (B) leafing trait, based on the best-supported result from the discrete model analyses. The pie chart at each node within the phylogenetic tree represents the proportional probability of the four states. The values below the phylogenetic tree indicate the relative transition rates among the four states. **(A)** Ancestral state reconstruction of the fruiting and pollination traits. The four states are represented by light pink, dark pink, light green, and dark green, corresponding to 1-year fruiting (1-year) & anemophily (AN), 1-year fruiting & entomophily (EN), 2-year fruiting (2-year) & anemophily, and 2-year fruiting & entomophily, respectively. **(B)** Ancestral state reconstruction of the fruiting and leafing traits. The four states are represented by light pink, dark pink, light green, and dark green, corresponding to 1-year fruiting & deciduous (DEC), 1-year fruiting & evergreen (EVE), 2-year fruiting & deciduous, and 2-year fruiting & evergreen, respectively.

## 4 Discussion

Our ancestral state reconstructions indicate that the 2-year fruiting originated once in the most recent common ancestor of the genera in which this trait is present today (node N3 uniting the six clades except *Fagus* and *Trigonobalanus*: see Figs. 2, S4A). Previous molecular dating studies have placed the emergence of this lineage in the late Cretaceous to early Paleocene (approximately 85–60 Ma; Liu et al., 2025; Yang et al., 2025; Zhou et al., 2022). Because this timeframe coincides with major global climate changes and the emergence of seasonality (Toumoulin et al. 2022), it is tempting to speculate that these environmental changes may have facilitated—or driven—the evolution of delayed fertilization and the resulting 2-year fruiting cycle. A more detailed understanding of this evolutionary context will require fossil-informed biogeographic analyses that explicitly link environmental dynamics with the origin and diversification of prolonged delayed fertilization.

Following the initial acquisition of the 2-year fruiting, multiple independent reversions to the 1-year fruiting were inferred across the family. These reversions occurred in the common ancestor of *Castanea*, within *Castanopsis*, and in several clades of *Quercus*, indicating that although shifts back to 1-year fruiting are relatively rare, they have arisen repeatedly across different genera. Additional cases are likely present in *Lithocarpus*, where several species not included in our analysis (e.g., *L. dodonaeifolius*, *L. formosanus*) are known to exhibit 1-year fruiting (Brach & Song, 2006). Notably, no cases of reversion of the 2-year fruiting were detected in any lineage that had reverted to the 1-year fruiting lineages. A prominent example involves the common ancestor of *Quercus* sect. *Ponticae*, sect. *Virentes*, and sect. *Quercus*, which arose during the Eocene (56–33.9 Ma). Although this ancestor is inferred to have undergone a transition back to the 1-year fruiting, its subsequent range expansion across Eurasia and North America was not accompanied by a secondary gain of the 2-year fruiting (Fig. 2; Fig. S4A; Denk et al. 2017; Hipp et al., 2018, 2020; Zhou et al., 2022). Likewise, in *Castanea*, all extant species are 1-year fruiting without any reversion to 2-year fruiting. Ancestral-state reconstructions strongly support entomophily and evergreen leaf habit as ancestral traits of Quercoideae (node N2; Figs. S4B, C) predating the emergence of the 2-year fruiting strategy at the most recent common ancestor (node N3). This pattern indicates that 2-year fruiting evolved against a background of entomophilous pollination and evergreen leaf habit. Insect pollination characterizes extant genera such as *Castanea*, *Castanopsis*, *Lithocarpus*, *Chrysolepis*, *Notholithocarpus*, and *Trigonobalanus verticillata*, which possess compound inflorescences and attract generalist insect pollinators (Chen et al. 2025; Crepet and Nixon 1989; Larue and Petit 2024; Larue et al. 2021; Kaul and Abbe 1984; Kaul 1986; Nixon and Crepet 1989; Petit and Larue 2022; Sadowski 2020; Soepadmo 1972; Wright and Dodd 2013; Yuan et al. 2024), whereas wind pollination in *Quercus* and *Trigonobalanus doichangensis* represents derived states (Manos et al. 2001; Nixon and Crepet 1989; Zhou et al. 2022); for *Fagus*, the ancestral or derived status of wind pollination remains unresolved, despite previous suggestions of wind pollination as ancestral in Fagales and Fagaceae (Stephens et al. 2023; Yao et al. 2023).

Evergreen leaf habit, likely acquired under warm Cretaceous–Paleocene climates, has been phylogenetically conserved in low-latitude Asian lineages such as *Castanopsis*, *Lithocarpus*, and some sections of *Quercus* (Sects. *Cyclobalanopsis* and *Ilex*), whereas independent transitions to deciduousness accompanied northward range expansion in *Castanea* and several *Quercus* lineages (Sects. *Lobatae* and *Quercus*), likely driven by colder and more seasonal climates (Barrón et al. 2017; Cavender-Bares 2018; Deng et al. 2018; Fontes et al. 2025; Hipp et al. 2018; Jiang et al. 2019; Kikuzawa 1991; Sancho-Knapik et al. 2021; Soepadmo 1972; Zanne et al. 2014). Multiple secondary transitions from deciduous to evergreen states further indicate that leaf habit evolution reflects both phylogenetic conservatism and repeated adaptive shifts (Hipp et al., 2018; Sancho-Knapik et al., 2021; Furze et al., 2021). Despite these pronounced evolutionary dynamics, our discrete-trait analyses detected no evidence for correlated evolution between fruiting strategy and either pollination mode or leaf habit. While the number of transitions inherently limits statistical power, this result suggests no strong coupling between these traits.

As genomic resources and transcriptome data for Fagaceae continue to expand (Satake et al., 2023; Kudo et al., 2025; Lara-De La Cruz and Chávez-Vergara, 2026), future studies aimed at uncovering the molecular and developmental bases of both the origin and reversion of 2-year fruiting will become increasingly feasible and highly informative. Genes governing ovule developmental timing and the environmental regulation of dormancy release represent promising targets (Satake et al., 2022; Satake et al., 2023). Integrating such molecular insights with comparative phylogenetic analyses will substantially advance our understanding of the evolutionary processes that gave rise to delayed fertilization in the Fagaceae.

## Supporting information

Suppementary Table 1

Suppementary Table 2

Suppementary Table 3

Suppementary Table 4

Suppementary Table 5

## Author contributions

**Takenori Shagawa:** data curation (lead), formal analysis (equal), methodology (equal), validation (equal), visualization (equal), writing—original draft (lead), writing—review and editing (equal). **Chihiro Myotoishi:** data curation (supporting), formal analysis (equal), methodology (equal), validation (equal), visualization (equal), writing—review and editing (equal). **Tetsukazu Yahara:** data curation (supporting), supervision (supporting), writing—review and editing (equal). **Ryosuke Imai:** methodology (equal), writing—review and editing (equal). **Min Deng:** data curation (supporting), writing—review and editing (equal). **Akiko Satake:** conceptualization (lead), funding acquisition (lead), methodology (equal), project administration (lead), supervision (lead), validation (equal), writing—review and editing (equal). Takenori Shagawa and Chihiro Myotoishi should be considered joint first authors.

## Acknowledgments

We are grateful to the curators of the following herbaria for providing access to online specimen data: A, ANDA, AU, BM, BR, CAF, CDBI, CSFI, E, GAC, GFS, GH, GZAC, HITBC, HK, IBK, IBSC, JXAU, K, KUN, L, MEL, MICH, MO, NAS, NY, P, PE, SHM, SN, SYS, SZ, US, and WU. We sincerely thank Ai Nagahama for her valuable discussions and insightful comments on this study. We gratefully acknowledge the Research Institute for Information Technology, Kyushu University, for providing the computational resources used for phylogenetic reconstruction under the General Projects category. This study was supported by JSPS KAKENHI (JP23H04965 and JP23H04966) awarded to A.S.

## Data Availability Statement

Trait data used in this study are publicly available from the eFloras of China (Brach and Song, 2006) and the eFlora of North America (Nixon, 1997). Additional literature sources used for trait data compilation are listed in Table S1. Information on digitized herbarium specimens used to verify and refine trait assignments is provided in Tables S2 and S3.

## Appendices

**FIGURE A1** Majority-rule consensus tree reconstructed with MrBayes v.3.2.7 from the nuclear genome dataset of Zhou et al. (2022), after removing excluded samples. Posterior probabilities (PP) are shown at each node. Scale bar indicates substitutions per site.

**FIGURE A2** Overview of the 52 alternative model settings considered in the multistate model analysis of fruiting traits. Model IDs are shown at the upper left of each tree, and branches assigned different transition rates are indicated in various colors.

**FIGURE A3** Schematic representation of four different model configurations in the discrete model analyses. In the independent model (left), each trait evolves separately with transition rates *α* and *β*, resulting in four parameters (unrestricted) or two parameters (restricted). In the dependent model (right), the two traits evolve jointly with transition rates *q*, resulting in eight parameters (unrestricted) or four parameters (restricted). Boxes indicate possible combined states of the two traits, and arrows represent allowed transitions with associated rate parameters.

**FIGURE A4** Estimated evolutionary history of the traits potentially associated with prolonged delayed fertilization, based on the best-supported results from the multistate model analyses. The pie chart at each node within the phylogenetic tree represents the proportional probability of alternative states, and the values below the phylogeny indicate the relative transition rates between states. **(A)** Ancestral state reconstruction of the fruiting trait when *Quercus suber* is treated as a 1-year fruiting species. Pink and green represent 1-year and 2-year fruiting, respectively. **(B)** Ancestral state reconstruction of the pollination trait. Light blue and yellow represent anemophily (AN) and entomophily (EN), respectively. **(C)** Ancestral state reconstruction of the leafing trait. Light brown and dark green represent deciduous (DEC) and evergreen (EVE), respectively.

## Supplementary Information

**TABLE S1** List of Fagaceae species analyzed and their trait information.

**TABLE S2** List of herbaria and corresponding data portals for cited Fagaceae specimens.

**TABLE S3** Online herbarium specimens used for data collection.

**TABLE S4** Summary of model settings and results of ancestral state reconstruction using BayesTraits. The model shown in bold was selected as the best-fitting model based on Bayes factor comparisons calculated from log marginal likelihoods.

**TABLE S5** Probability of trait states at nodes N1–N14 under the best[fit multistate models for each analyzed trait dataset.

## References

1. Akaike, H. 1974. “A new look at the statistical model identification.” IEEE Transactions on Automatic Control 19(6): 716–723. 10.1109/TAC.1974.1100705

2. Araye, Q., Yahara, T., and Satake, A. 2023. “Latitudinal cline of flowering and fruiting phenology in Fagaceae in Asia.” Biotropica 55(1): 277–285. 10.1111/btp.13184

3. Barrón, E., Averyanova, A., Kvacek, Z., Momohara, A., Pigg, K. B., Popova, S., Postigo-Mijarra, J. M., Tiffney, B. H., Utescher, T., and Zhou, Z. K. 2017. “The fossil history of *Quercus*.” In: Gil-Pelegrın, E., Peguero-Pina, J. J., Sancho-Knapik, D. (Eds.) Oaks physiological ecology. Exploring the functional diversity of genus Quercus L.: 39–105. Cham, Switzerland, Springer.

4. Benson, M. 1894. “Contribution to the embryology of the *Amentiferae*. Part I.” Transactions of the Linnean Society of London. 2nd Series. Botany 3(10): 409–424. 10.1111/j.1095-8339.1894.tb00624.x

5. Brach, A. R., and Song, H. 2006. “eFloras: New directions for online floras exemplified by the flora of China project.” Taxon 55(1): 188–192. 10.2307/25065540

6. Cavender-Bares, J. 2018. “Diversification, adaptation, and community assembly of the American oaks (*Quercus*), a model clade for integrating ecology and evolution.” New Phytologist 221(2): 669–692. 10.1111/nph.15450

7. Chen, X., Wang, H., Ren, Z.X., and Liu, P. 2025. “Stone oak flowers are more likely generalist pollinated by insects rather than wind.” Plant Ecology 226(8): 933–943. 10.1007/s11258-025-01541-x

8. Crepet, W. L., and Nixon, K. C. 1989. “Earliest megafossil evidence of Fagaceae: phylogenetic and biogeographic implications.” American Journal of Botany 76(6): 842–855. 10.1002/j.1537-2197.1989.tb15062.x

9. Deng, M., Jiang, X. L., Hipp, A. L., Manos, P. S., and Hahn, M. 2018. “Phylogeny and biogeography of East Asian evergreen oaks (*Quercus* section *Cyclobalanopsis*; Fagaceae): Insights into the Cenozoic history of evergreen broad-leaved forests in subtropical Asia.” Molecular Phylogenetics and Evolution 119: 170–181. 10.1016/j.ympev.2017.11.003

10. Deng, M., Yao, K., Shi, C., Shao, W., and Li, Q. 2022. “Development of *Quercus acutissima* (Fagaceae) pollen tubes inside pistils during the sexual reproduction process.” Planta 256(1): 16. 10.1007/s00425-022-03937-9

11. Denk, T., Grimm, G. W., Manos, P. S., Deng, M., and Hipp, A. L. 2017. “An updated infrageneric classification of the oaks: review of previous taxonomic schemes and synthesis of evolutionary patterns.” In: Gil-Pelegrın, E., Peguero-Pina, J. J., Sancho-Knapik, D. (Eds.) Oaks physiological ecology. Exploring the functional diversity of genus Quercus L.: 13–38. Cham, Switzerland, Springer.

12. Díaz-Fernández, P. M., Climent, J., and Gil, L. 2004. “Biennial acorn maturation and its relationship with flowering phenology in Iberian populations of *Quercus suber*.” Trees 18(6): 615–621. 10.1007/s00468-004-0325-z

13. Elena-Rossello, J. A., de Rio, J. M., Valdecantos J. L. G., and Santamaria I. G. 1993. “Ecological aspects of the floral phenology of the cork-oak (*Q suber* L): why do annual and biennial biotypes appear?” Annals of Forest Science 50(Suppl.): 114s–121s. 10.1051/forest:19930710Ann

14. Fontes, C. G., Meireles, J. E., Hipp, A. L., and Cavender-Bares, J. 2025. “Adaptive evolution of freezing tolerance in oaks is key to their dominance in North America.” Ecology Letters 28(2): e70084. 10.1111/ele.70084

15. Furze, M. E., Wainwright, D. K., Huggett, B. A., Knipfer, T., McElrone, A. J., and Brodersen, C. R. 2021. “Ecologically driven selection of nonstructural carbohydrate storage in oak trees.” New Phytologist 232(2): 567–578. 10.1111/nph.17605

16. Givnish, T. J. 2002. “Adaptive significance of evergreen vs. deciduous leaves: solving the triple paradox.” Silva fennica 36(3): 703–743. 10.14214/sf.535

17. Hipp, A. L., Manos, P. S., González-Rodríguez, A., Hahn, M., Kaproth, M., McVay, J. D., Valencia-Avalos, S., and Cavender-Bares, J. 2018. “Sympatric parallel diversification of major oak clades in the Americas and the origins of Mexican species diversity.” New Phytologist 217(1): 439–452. 10.1111/nph.14773

18. Hipp, A. L., Manos, P. S., Hahn, M., Avishai, M., Bodénès, C., Cavender-Bares, J., Crowl, A. A., Deng, M., Denk, T., Fitz-Gibbon, S., Gailing, O., González-Elizondo, M. S., González-Rodríguez, A., Grimm, G. W., Jiang, X. L., Kremer, A., Lesur, I., McVay, J. D., Plomion, C., Rodríguez-Correa, H., Schulze, E. D., Simeone, M. C., Sork, V. L., and Valencia-Avalos S. 2020. “Genomic landscape of the global oak phylogeny.” New Phytologist 226(4): 1198–1212. 10.1111/nph.16162

19. Jiang, X. L., Hipp, A. L., Deng, M., Su, T., Zhou, Z. K., and Yan, M. X. 2019. “East Asian origins of European holly oaks (*Quercus* section *Ilex* Loudon) via the Tibet-Himalaya.” Journal of Biogeography 46(10): 2203–2214. 10.1111/jbi.13654

20. Kass, R. E., and Raftery, A. E. 1995. “Bayes Factors.” Journal of the American Statistical Association 90(430): 773–795. 10.2307/2291091

21. Kaul, R. B. and Abbe, E. C. 1984. “Inflorescence architecture and evolution in the Fagaceae.” Journal of the Arnold Arboretum 65(3): 375–401. https://www.jstor.org/stable/43782573

22. Kaul, R. B. 1986. “Evolution and reproductive biology of inflorescences in *Lithocarpus*, *Castanopsis*, *Castanea*, and *Quercus* (Fagaceae).” Annals of the Missouri Botanical Garden 73(2): 284–296. 10.2307/2399114

23. Kikuzawa, K. 1991. “A cost-benefit analysis of leaf habit and leaf longevity of trees and their geographical pattern.” The American Naturalist 138(5): 1250–1263. 10.1086/285281

24. Kudo, S. N., Ikezaki, Y., Kusumi, J., Hirakawa, H., Isobe, S., and Satake, A. 2025. “The evolution of gene expression in seasonal environments.” eLife 14: RP107309. 10.7554/eLife.107309

25. Lanfear, R., Frandsen, P. B., Wright, A. M., Senfeld, T., and Calcott, B. 2017. “PartitionFinder 2: new methods for selecting partitioned models of evolution for molecular and morphological phylogenetic analyses.” Molecular Biology and Evolution 34(3): 772–773. 10.1093/molbev/msw260

26. Lara-De La Cruz, L. I., and Chávez-Vergara, B. 2026. “Transcriptomics in oaks (Fagaceae; *Quercus*): A comprehensive review of advances, biases, and future directions.” Botanical Sciences 104(1): (early view). 10.17129/botsci.3717

27. Larue, C., Austruy, E., Basset, G., and Petit, R. J. 2021. “Revisiting pollination mode in chestnut (*Castanea* spp.): an integrated approach.” Botany Letters 168(3): 348–372. 10.1080/23818107.2021.1872041

28. Larue, C., and Petit, R. J. 2024. “Insect pollination in chestnut: an organized mess?” Acta Horticulturae 1400: 331–340. 10.17660/ActaHortic.2024.1400.40

29. Liu, S. Y., Yang, Y. Y., Tian, Q., Yang, Z. Y., Li, S. F., Valdes, P. J., Farnsworth, A., Kates, H. R., Siniscalchi, C. M., Guralnick, R. P., Soltis, D. E., Soltis, P. S., Stull, G. W., Folk, R. A., and Yi, T. S. 2025. An integrative framework reveals widespread gene flow during the early radiation of oaks and relatives in Quercoideae (Fagaceae). Journal of Integrative Plant Biology 67(4): 1119–1141. 10.1111/jipb.13773

30. Manos, P. S., Zhou Z. K., and Cannon, C. H. 2001. “Systematics of Fagaceae: phylogenetic tests of reproductive trait evolution.” International Journal of Plant Sciences 162(6): 1361–1379. 10.1086/322949

31. Nixon, K. C. 1997. “Fagaceae.” In: Flora of North America Editorial Committee (Eds.), Flora of North America 3. Oxford University Press, New York. 436–506. http://floranorthamerica.org/Fagaceae [2025.02.06 accessed.]

32. Nixon, K. C., and Crepet, W. L. 1989. “*Trigonobalanus* (Fagaceae): taxonomic status and phylogenetic relationships.” American Journal of Botany 76(6): 828–841. 10.1002/j.1537-2197.1989.tb15061.x

33. Pagel, M. 1994. “Detecting correlated evolution on phylogenies: a general method for the comparative analysis of discrete characters.” Proceedings of the Royal Society B: Biological Sciences 255: 37–45. 10.1098/rspb.1994.0006

34. Pagel, M., Meade, A., and Barker, D. 2004. “Bayesian Estimation of Ancestral Character States on Phylogenies.” Systematic Biology 53(5): 673–684. 10.1080/10635150490522232

35. Pagel, M., and Meade, A. 2006. “Bayesian Analysis of Correlated Evolution of Discrete Characters by Reversible[Jump Markov Chain Monte Carlo.” The American Naturalist 167(6): 808–825. 10.1086/503444

36. Paradis, E., Claude, J., and Strimmer, K. 2004. “APE: Analyses of Phylogenetics and Evolution in R language.” Bioinformatics 20(2): 289–290. 10.1093/bioinformatics/btg412

37. Paradis, E., and Schliep, K. 2019. “ape 5.0: an environment for modern phylogenetics and evolutionary analyses in R.” Bioinformatics 35(3): 526–528. 10.1093/bioinformatics/bty633

38. Petit, R. J., and Larue, C. 2022. “Confirmation that chestnuts are insect-pollinated.” Botany Letters 169(3): 370–374. 10.1080/23818107.2022.2088612

39. Ronquist, F., Teslenko, M., van der Mark, P., Ayres, D. L., Darling, A., Höhna, S., Larget, B., Liu, L., Suchard, M. A., and Huelsenbeck, J. P. 2012. “MrBayes 3.2: Efficient bayesian phylogenetic inference and model choice across a large model space.” Systematic Biology 61(3): 539–542. 10.1093/sysbio/sys029

40. Sadowski, E. M., Schmidt, A. R., and Denk, T. 2020. “Staminate inflorescences with in situ pollen from Eocene Baltic amber reveal high diversity in Fagaceae (oak family).” Willdenowia 50(3): 405–517. 10.3372/wi.50.50303

41. Sancho-Knapik, D., Escudero, A., Mediavilla, S., Scoffoni, C., Zailaa, J., Cavender-Bares, J., Álvarez-Arenas, T. G., Molins, A., Alonso-Forn, D., Ferrio, J. P., Peguero-Pina, J. J., and Gil-Pelegrín E. 2021. “Deciduous and evergreen oaks show contrasting adaptive responses in leaf mass per area across environments.” New Phytologist 230(2): 521–534. 10.1111/nph.17151

42. Satake, A., Nagahama, A., and Sasaki, E. 2022. “A cross-scale approach to unravel the molecular basis of plant phenology in temperate and tropical climates.” New Phytologist 233(6): 2340–2353. 10.1111/nph.17897

43. Satake, A., Ohta, K., Takeda-Kamiya, N., Toyooka, K., and Kusumi, J. 2023. “Seasonal gene expression signatures of delayed fertilization in Fagaceae.” Molecular Ecology 32(17): 4801–4813. 10.1111/mec.17079

44. Satake, A., and Kelly, D. 2021. “Delayed fertilization facilitates flowering time diversity in Fagaceae.” Philosophical Transactions of the Royal Society B 376(1839): 20210115. 10.1098/rstb.2021.0115

45. Schoonderwoerd, K. M., and Friedman, W. E. 2016. “Zygotic dormancy underlies prolonged seed development in *Franklinia alatamaha* (Theaceae): a most unusual case of reproductive phenology in angiosperms.” Botanical Journal of the Linnean Society 181(1): 70–83. 10.1111/boj.12409

46. Schwartz, M. D. 2003. Phenology: an integrative environmental science. Springer.

47. Shagawa, T., Ogawa, K., Kanaoka, M. M., and Satake, A. 2025. “A seasonal strategy for pollen tube growth and ovule development to overcome winter in Japanese stone oak (*Lithocarpus edulis*).” Scientific Reports 15: 16131. 10.1038/s41598-025-00529-x

48. Shen, Z., Zhou, B. F., Liang, Y. Y., Wang, J. S., Yu, R. X., Shi, Y., Ling, S. J., Luo, W. J., Lin, Q. Q., Niu, J. W., Qiao, L. J., Manos, P. S., and Wang, B. 2025. “Teasing apart the sources of phylogenetic tree discordance across three genomes in the oak family (Fagaceae).” BMC Plant Biology 25: 919. 10.1186/s12870-025-06963-3

49. Soepadmo, E. 1972. “Fagaceae.” Flora Malesiana Series 1. Spermatophyta. 7: 265–403.

50. Sogo, A., and Tobe, H. 2006. “Delayed fertilization and pollen-tube growth in pistils of *Fagus japonica* (Fagaceae).” American Journal of Botany 93(12): 1748–1756. 10.3732/ajb.93.12.1748

51. Stephens, R. E., Gallagher, R. V., Dun, L., Cornwell, W., and Sauquet, H. 2023. “Insect pollination for most of angiosperm evolutionary history.” New Phytologist 240(2): 880–891. 10.1111/nph.18993

52. Toumoulin, A., Tardif, D., Donnadieu, Y., Licht, A., Ladant, J. B., Kunzmann, L., and Dupont-Nivet, G. 2022. “Evolution of continental temperature seasonality from the Eocene greenhouse to the Oligocene icehouse –a model–data comparison.” Climate of The Past 18(2): 341–362. 10.5194/cp-18-341-2022

53. Williams, J. H. 2008. “Novelties of the flowering plant pollen tube underlie diversification of a key life history stage.” PNAS 105(32): 11259–11263. 10.1073/pnas.0800036105

54. Williams, J. H., and Reese, J. B. 2019. “Evolution of development of pollen performance.” Current Topics in Developmental Biology 131: 299–336. 10.1016/bs.ctdb.2018.11.012

55. Wright, J. W., and Dodd, R. S. 2013. “Could tanoak mortality affect insect biodiversity? Evidence for insect pollination in tanoaks.” Madroño 60(2): 87–94. 10.3120/0024-9637-60.2.87

56. Xie, W., Lewis, P. O., Fan, Y., Kuo, L., and Chen, M. H. 2011. “Improving marginal likelihood estimation for Bayesian phylogenetic model selection.” Systematic Biology 60(2): 150–160. 10.1093/sysbio/syq085

57. Yang, Y., Y., Stull, G. W., Qu, X. J., Deng, M., Zhao, L., Hu, Y., Wang, Z. H., Ma, H., Li, D. Z., Smith, S. A., and Yi, T. S. 2025. “Genome duplications, genomic conflict, and rapid phenotypic evolution characterize the Cretaceous radiation of Fagales.” Journal of Integrative Plant Biology 67(11): 2929–2944. 10.1111/jipb.70011

58. Yao, K., Deng, M., Lin, L., Hu, J., Yang, X., Li, Q., and Feng, Z. 2023. “The fertilization process in *Lithocarpus dealbatus* (Fagaceae) and its implication on the sexual reproduction evolution of Fagales.” Planta 258(2): 23. 10.1007/s00425-023-04178-0

59. Yuan, B., Li, Y., Zhang, J., Zhang, X., Hu, F., Yuan, D., and Fan, X. 2024. “Managing flower-visiting insects is essential in *Castanea*: Enhance yield while ensuring quality.” iScience 27(11): 111127. 10.1016/j.isci.2024.111127

60. Zanne, A. E., Tank, D. C., Cornwell, W. K., Eastman, J. M., Smith, S. A., FitzJohn, R. G., McGlinn, D. J., O’Meara, B. C., Moles, A. T., Reich, P. B., Royer, D. L., Soltis, D. E., Stevens, P. F., Westoby, M., Wright, I. J., Aarssen, L., Bertin, R. I., Calaminus, A., Govaerts, R., Hemmings, F., Leishman, M. R., Oleksyn, J., Soltis, P. S., Swenson, N. G., Warman, L., and Beaulieu, J. M. 2014. “Three keys to the radiation of angiosperms into freezing environments.” Nature 506: 89–92. 10.1038/nature12872

61. Zhou, B. F., Yuan, S., Crowl, A. A., Liang, Y. Y., Shi, Y., Chen, X. Y., An, Q. Q., Kang, M., Manos, P. S, and Wang B. 2022. “Phylogenomic analyses highlight innovation and introgression in the continental radiations of Fagaceae across the northern hemisphere.” Nature Communications 13: 1320. 10.1038/s41467-022-28917-1

