## Supplementary figures and images for "Evolutionary origin of prolonged delayed fertilization in the Fagaceae"

### Appendix 1

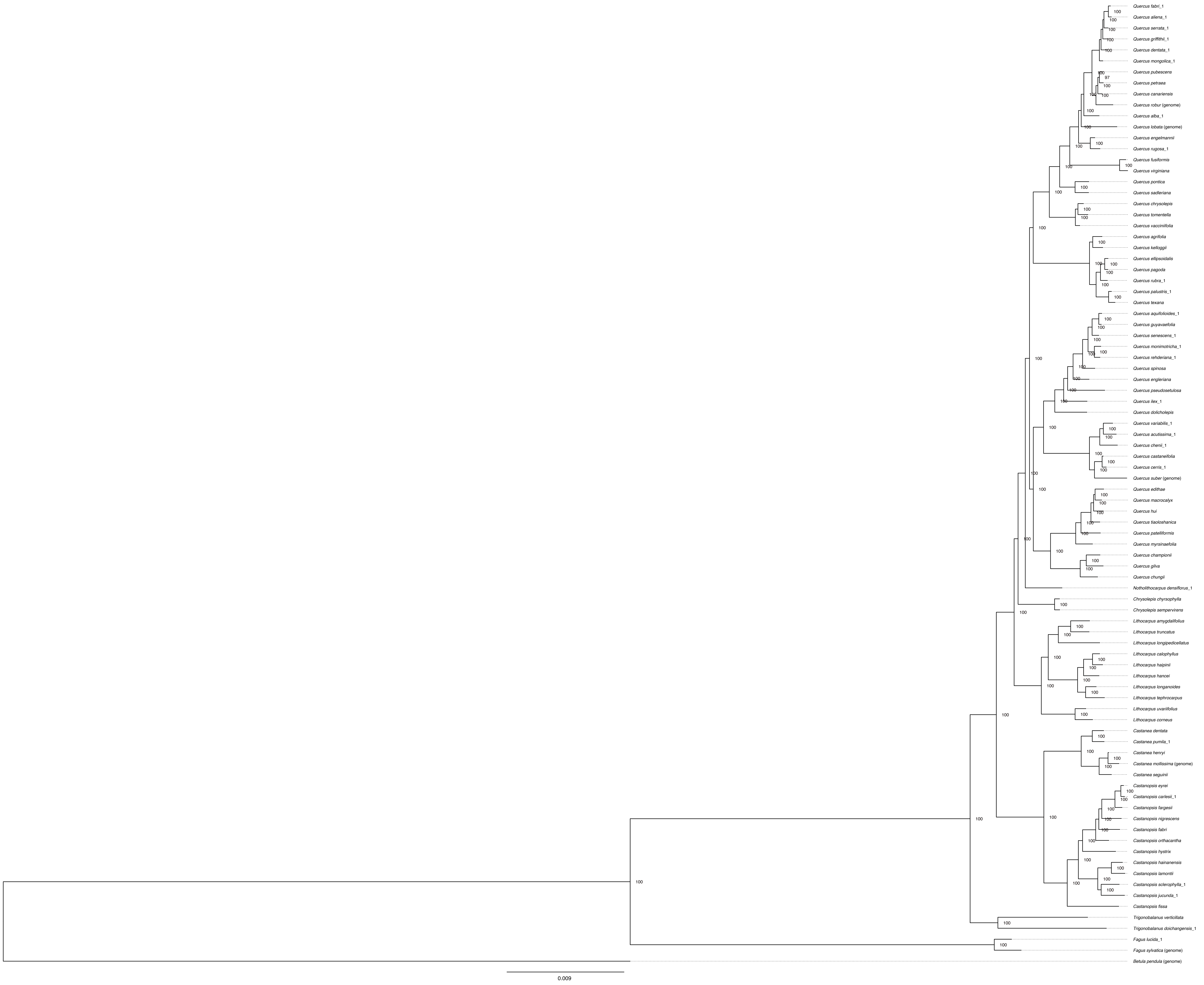

### Appendix 2

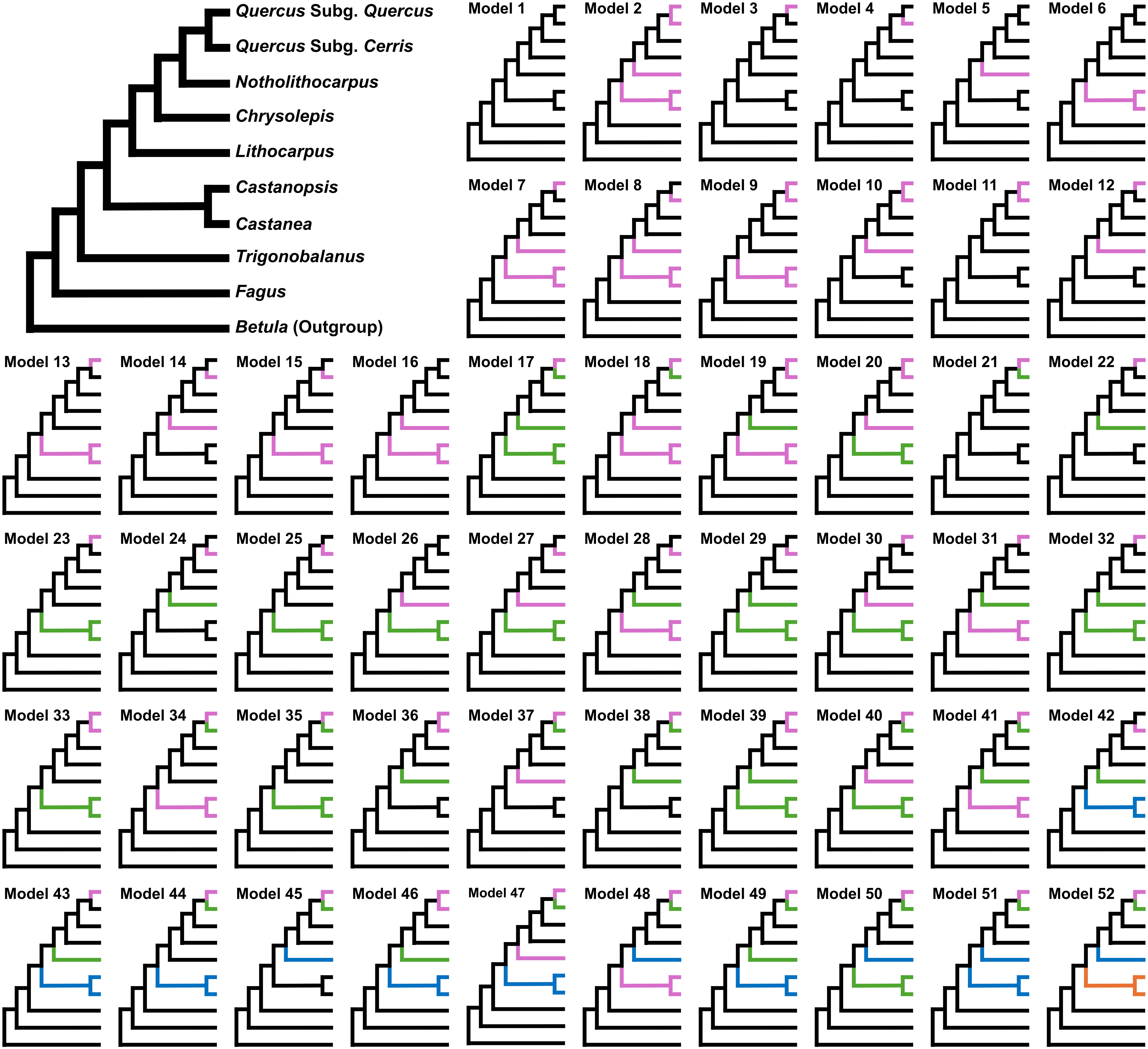

### Appendix 3

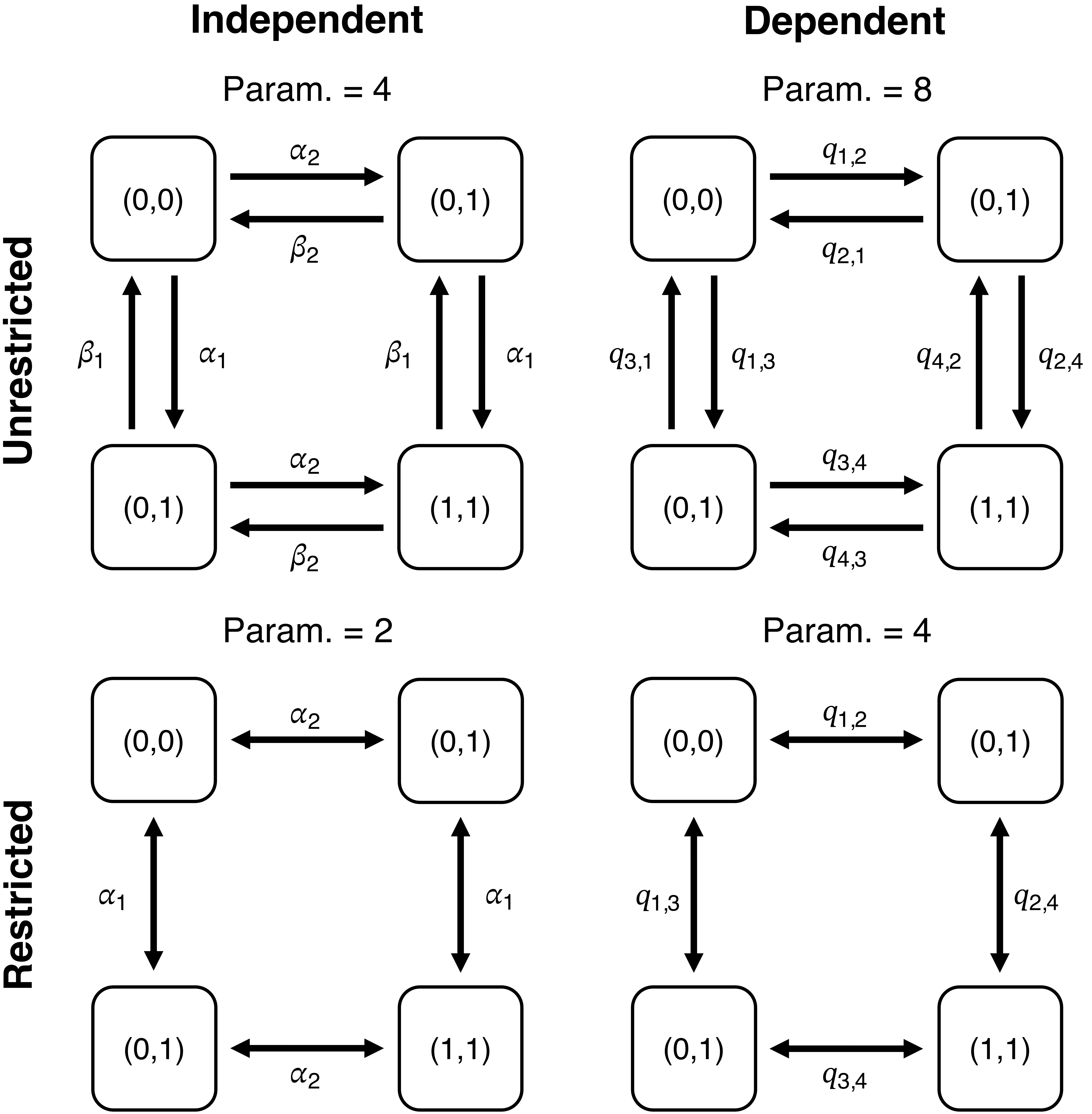

### Appendix 4

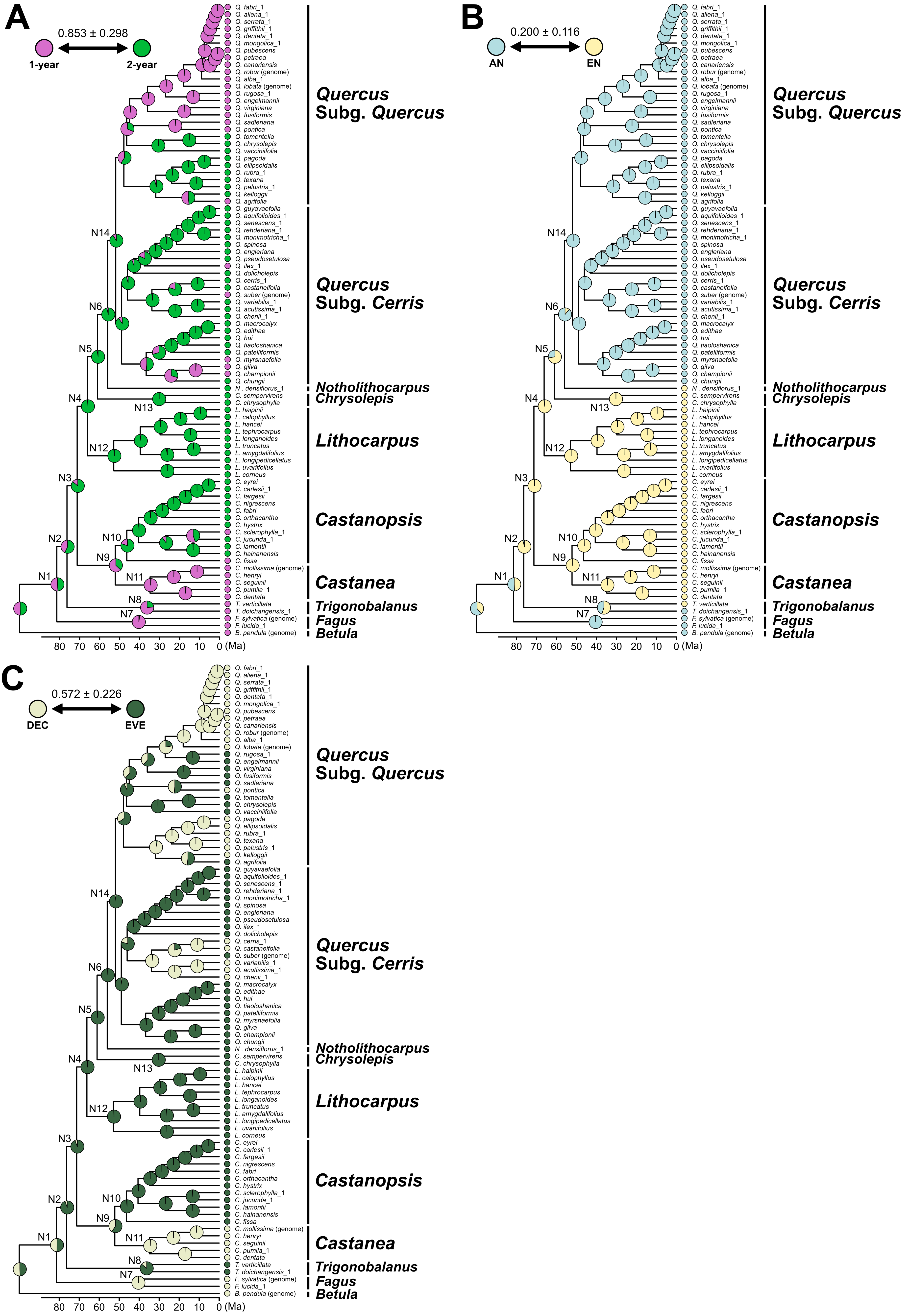
